# The structure of plant spatial association networks increases plant diversity in global drylands

**DOI:** 10.1101/121491

**Authors:** H. Saiz, J. Gómez-Gardeñes, J.P. Borda, F.T. Maestre

**Affiliations:** Departamento de Biología y Geología, Física y Química Inorgánica, Universidad Rey Juan Carlos. C/ Tulipán s/n, 28933 Móstoles, SPAIN.; Departamento de Física de la Materia Condensada, Universidad de Zaragoza. C/ Pedro Cerbuna 12, 50009 Zaragoza, SPAIN.; Institute for Biocomputation and Physics of Complex Systems (BIFI), Universidad de Zaragoza. C/ Mariano Esquillor (Edificio I+D), 50018, Zaragoza, SPAIN.

**Keywords:** Competition, Drylands, Ecological networks, Facilitation, Plant diversity, Signed networks, Spatial patterns

## Abstract

**Aim:** Despite their widespread use and value to unveil the complex structure of the interactions within ecological communities and their value to assess the resilience of communities, network analyses have seldom been applied in plant communities. We aim to evaluate how plant-plant interaction networks vary in global drylands, and to assess whether network structure is related to plant diversity in these ecosystems.

**Location:** 185 dryland ecosystems from all continents except Antarctica.

**Methods:** We built networks using the local spatial association between all the perennial plant species present in the communities studied, and used structural equation models to evaluate the effect of abiotic factors (including geography, topography, climate and soil conditions) and network structure on plant diversity.

**Results:** The structure of plant networks found at most study sites (72%) was not random and presented properties representative of robust systems, such as high link density and structural balance. Moreover, network indices linked to system robustness had a positive and significant effect on plant diversity, sometimes higher that the effect of abiotic factors.

**Main conclusions:** Our results constitute the first empirical evidence showing the existence of a common network architecture structuring terrestrial plant communities at the global scale, and provide novel evidence of the importance of the network of interactions for the maintenance of biodiversity. Furthermore, they highlight the importance of system-level approaches to explain the diversity and structure of interactions in plant communities, two major drivers of terrestrial ecosystem functioning and resilience against the likely impacts derived from global change.

## Introduction

Network analyses are being increasingly used in ecology to unveil the complexity of species interactions and to study their effects on the functioning and stability of ecosystems (Ings *et al*., 2009; Heleno *et al*., 2014). Theoretical studies have linked particular network properties with the stability of ecological communities (Rohr *et al*., 2014; Allesina *et al*., 2015), and it has been hypothesized that real ecological networks are more prone to have properties promoting the efficiency of ecosystem processes (*e.g*. nutrient uptake, Arditi *et al*., 2005) and the robustness of communities against perturbations (Estrada, 2006). However, most studies in ecological networks have focused in a few specific systems (*e.g*. food webs, plant-pollinator, host-parasite) and have been conducted at particular study sites, making the establishing of generalizations difficult (Heleno *et al*., 2014). Thus, comparative studies of networks at regional and global scales are necessary to evaluate whether ecological communities present common properties across multiple environmental conditions, and to explore how they affect key ecosystem attributes such as species diversity and ecosystem functioning (Traveset *et al*., 2016).

Plant communities are the bottom of the trophic web, play a major role in ecosystem nutrient cycling and are responsible of community physiognomy (Barbour, 1987). Despite their critical ecological role, and the long tradition of ecological studies with plants, they have been largely unnoticed by network studies until very recently (Verdú *et al*., 2008; Saiz *et al*., 2016). The efforts required for obtaining data on plant-plant interactions at community level over a large number of sites (Soliveres & Maestre, 2014), and the different interactions that can be established between plants, such as facilitation or competition (Brooker *et al*., 2008), have traditionally hampered the use of network analyses to study the structure of plant communities. However, these limitations are starting to be overcome with the increase in the number of coordinated experiments and surveys being conducted globally (Maestre *et al*., 2012b; Fraser *et al*., 2013), and with methodological developments in the analysis of social networks with positive and negative links (*e.g*. like and dislikes; (Doreian & Mrvar, 2009; Szell *et al*., 2010). To our knowledge, no study so far has evaluated the structure of plant networks and how it relates to the diversity of plant communities at the global scale. Such analyses would help to unveil global patterns for plant communities, providing simultaneously information about the community (*e.g*. the relative importance of positive and negative interactions) and the species forming it (*e.g*. the role of each species structuring the community, Saiz *et al*., 2014). Furthermore, the connection between the structure and resilience against extinctions of the network will provide a valuable information about the vulnerability of plant communities facing possible future extinctions due to global change.

In this study we explored the structure of plant spatial networks, and evaluated its effects on the diversity of plant communities in global drylands. Despite covering over 45% of global terrestrial area (Prăvălie, 2016), few studies so far have evaluated the network structure of dryland plant communities (Saiz & Alados, 2014; Saiz *et al*., 2014). Understanding the network structure of dryland plant communities is particularly relevant for multiple reasons. Dryland vegetation is organized as patches embedded in a matrix of bare soil, which become sinks for resources (*e.g*. rainfall, Aguiar & Sala, 1999; Wang *et al*., 2007). Species responsible of patch formation (‘nurses’) create a microenvironment where other species, less tolerant to dry environmental conditions, are able to establish (Maestre *et al*., 2001). Thus, positive interactions play a key role structuring plant communities in drylands, and allow the persistence of communities with higher biodiversity (Verdú *et al*., 2008; Soliveres & Maestre, 2014). However, a network approach can include not only facilitation, but also negative interactions, which also are important drivers in structuring dryland plant communities (Fowler, 1986; Soliveres *et al*., 2015b). Moreover, several studies have linked vegetation patchiness with ecosystem processes (Berdugo *et al*., 2017), so we could expect that plant network structure will have a direct effect on the functioning of dryland ecosystems. On the other hand, it has been predicted that drylands will experience the greatest proportional change in biodiversity in the near future (Sala *et al*., 2000), so a better understanding of the connection between the plant-plant interaction network structure and plant diversity can provide clues about how dryland communities will respond to the upcoming changes that they will face under ongoing global environmental change.

Dryland vegetation presents a marked spatial organization resulting from the interplay between climatic conditions, soil properties and plant-plant interactions (Sala & Aguiar, 1996; Rietkerk *et al*., 2004). Therefore, we expect that, under a given soil conditions and climate, this organization will be a good proxy of the structure of interactions between plants. Hence, we built plant-plant networks with positive and negative links considering the spatial association between all perennial plants present in 185 plant communities from all continents except Antarctica, and used structural equation models (Grace, 2006) for testing the effect of network structure on the plant community diversity. Specifically, we hypothesize that in drylands a) plant spatial networks present a common structure that promotes the resilience of the system against the extinction of species and interactions, and b) this structure has a direct effect on the diversity of the plant community. We expect that plant spatial networks in drylands present a high number and variety of connections between species (*i.e*. high link density and heterogeneity) due to the importance of biotic interactions, and to that of facilitation in particular, as drivers of community assembly. Furthermore, and after controlling the effect of abiotic factors, we anticipate that both a high link density and heterogeneity and the dominance of positive links will have a significant and positive effect on the diversity of dryland plant communities.

## Methods

### Global drylands vegetation survey

Field data were collected from 185 dryland sites located in 17 countries (Argentina, Australia, Botswana, Brazil, Burkina Faso, Chile, China, Ecuador, Ghana, Iran, Kenya, Mexico, Morocco, Peru, Spain, Tunisia, USA and Venezuela). These sites are a subset of the global network of 236 sites from Ulrich *et al*. (2016). As network indices depend on network size and must be tested against null models (Dormann *et al*., 2009), we selected all sites which networks had at least 5 connected species to allow statistical testing (185 sites out of 236 sites available). This subset included the major vegetation types found in drylands, a wide range in plant species richness (from 5 to 52 species per site) and environmental conditions (mean annual temperature and precipitation ranged from -1.8 to 28.2 <u>°</u>C, and from 66 to 1219 mm, respectively).

At each site, vegetation was surveyed using four 30-m-long transects located parallel among them within a 30 m × 30 m plot representative of the vegetation found there (see Maestre *et al*., 2012b for details). At each transect, 20 quadrats of 1.5 m × 1.5 m were established, and the cover of each perennial species within each quadrat was visually estimated

### Network construction

For each of the study sites, we built a plant-plant spatial association network (Saiz *et al*., 2014, 2016) using the cover data of all the perennial species (*S*) surveyed. These networks are characterized by the adjacency graph **A**_**S×S**_ (hereafter **A**), where the nodes (*i,j*) are the plant species and the links (*l*_*ij*_) are the spatial association between each pair of species. To determine this association, we calculated the correlation between the cover of each pair of species in the quadrats within each site using Spearman rank tests. When a correlation between species *i* and *j* was significant (*p* < 0.05), a link *l*_*ij*_ = *ρ* was established (where *ρ* represents the Spearman correlation coefficient), with *l_ij_* = 0 otherwise. Thus, links in our networks are signed (can have positive and negative values) and weighted (can present values between −1 and +1). For each site, *L* represented the total number of links in the network. As each species only had a single cover value at each quadrat, we could not evaluate the intra-specific spatial association; thus, we set the diagonal of **A** to zero.

We are aware that the use of spatial association to infer real biotic interactions presents several limitations. For example, factors different from biotic interactions, such as plant dispersal strategies or environmental heterogeneity, can influence vegetation spatial patterns (Escudero *et al*., 2005; Getzin *et al*., 2008). However, it has been suggested that biotic interactions are the main driver of spatial pattern formation at local scales (Pearson & Dawson, 2003; Morales-Castilla *et al*., 2015), and previous studies have found a good agreement between the outcome of spatial association and that of experiments evaluating plant-plant interactions in dryland communities (Tirado & Pugnaire, 2003). Therefore, we considered spatial pattern in a local scale, similar to that used in other studies about biotic interactions in plant communities (Cavieres *et al*., 2006; Verdú & Valiente-Banuet, 2011). Another possible shortcoming of our approach is that different types of biotic interactions can lead to the same spatial pattern, like facilitation and parasitism (*i.e*. positive spatial association). Thus, it would not be possible to differentiate between both interactions. However, we did not find parasitic species in our sites, so we can approach that positive spatial association only represents facilitative interactions.

### Network indices

We selected four network indices to characterize the structure of the communities studied: link density, link weight mean, link weight heterogeneity, and global network balance. Link density (D) is the average number of links per node in the network (*D* = *L*/*S*), and represents the importance of spatial patterns in the plant community, with high *D* describing a community where vegetation is more spatially structured (*i.e*. significant spatial association between pairs of species are common). Link weight mean 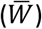 is the mean of all link weights in the network 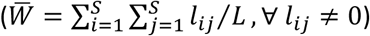 and represents the dominant type of links in the network, with 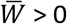 and 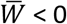 describing a community dominated by spatial aggregation and segregation, respectively. Link weight heterogeneity (*H*) is the kurtosis of the link weight distribution, with high *H* indicating a network where most links present similar weights. Global network balance (*K*) is a specific index for signed networks that accounts for the proportion of closed cycles in the network fulfilling the structural balance criterion (Zaslavsky, 2013). Following this criterion, a network can be divided in blocks; nodes within the same block are positively connected among them while they are negatively connected to nodes in other blocks (Doreian & Mrvar, 2009; Traag & Bruggeman, 2009). We calculated *K* using the definition of Estrada & Benzi (2014), 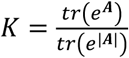 where |**A**| is the underlying unsigned graph of A. High values of *K* indicate that the network presents a ‘balanced’ structure (with *K* = 1 indicating a perfect balance), while low values indicate that several links do not fulfill this criterion and network is ‘unbalanced’ (*sensu*, network is ‘frustrated’, Doreian & Mrvar, 2009).

### Null model analyses

To test the significance of the network indices used, we employed two different null models for each network: one that allowed changing the connectivity of the network, and another that allowed changing the links between nodes while keeping the network linkage distribution constant. In the first model, we randomized the cover of each species along the quadrats. Specifically, we kept the cover distribution for each species constant, but changed randomly their positions in the quadrats. This way, we changed the cover values of species co-occurring in the same quadrat while maintaining the original cover distribution for each species at each site. Then, we built a network using this simulated data and calculated its *D*, 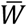 and H. For each site, we simulated 2000 networks and compared the real values of the indices against a 95% confidence interval created from the simulated networks. In the second null model, we simulated networks at each site using an algorithm based on the configurational model adapted for signed networks (Saiz *et al*., 2016). This method iteratively changes links in the original network, modifying its structure but keeping constant its linkage distribution. In our case, we made 1000 iterations per network and simulated 1000 networks, and we calculated *K* for each of the simulated networks. We also calculated the maximal and minimal *K* (*K*_*max*_ and *K*_*min*_) that each network could have considering its degree distribution to evaluate the real *K* value against all the possible values that it could present at each site. To do so, we iteratively simulated networks with the same null model, and selected the network that maximized (or minimized) *K* at each step. To avoid possible local maxima (or minima), selection was based on a Fermi-Dirac probability function 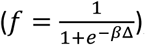, which selected a network over others based on the difference between the *K* values of the networks (Δ) and a parameter *β* that modulates the probability of accepting a change with the number of iterations (with higher *β* selecting higher Δ, Tsallis & Stariolo, 1996). By doing so we could precisely locate real networks in all the space of parameters of *K*.

### Evaluating the effects of network structure on plant diversity

We built structural equation models (SEM, Grace, 2006) including different abiotic variables (Latitude, Longitude, Elevation, Slope, Aridity, Seasonality, Soil organic C, Soil pH and soil total P) and network variables as explanatory variables for the richness (*SR*) and evenness (*E*) of perennial plant species. Specifically, we divided abiotic variables in four groups: geographical, topographical, climatic, and soil variables, and created a composite variable for each of them. We included the difference between real network values and the percentile 50 values for the networks simulated with the null models (*e.g*. Δ*D* = *D* − *D*_*null*_, where *D*_*null*_ is the percentile 50 for the *D* simulated with the null model) to remove random effects due to species abundance distribution (Gotelli, 2000) and network size (Dormann *et al*., 2009) from network indices. We then created a SEM for each combination of network and diversity variables (for a total of 4 × 2 = 8 SEMs). In these SEMs, network variables depended on all the composite variables, and diversity indices depended on all the composite variables and the network indices (see supplementary figures S1 and S2 for a complete description of the variables and the structure of the SEMs used). Specific dependencies between composite variables were included following previous studies using the same dataset (Delgado-Baquerizo *et al*., 2016). All variables were centered and standardized before calculating the models. All the SEM analyses were performed with the *lavaan* package (Rosseel, 2012) for R.3.2.4 (R Development Core TeamTeam, 2014).

## Results

The analysis of plant spatial association networks revealed that dryland plant communities present a wide variety of linkage structures (*i.e*. network indices were quite variable, Table 1, Figure 1). Particularly, 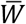 presented both positive and negative values, suggesting that drylands can be dominated by spatial aggregation or segregation. However, *K* presented a very low variability, with values close to 1 (Table 1). These results suggested that, in general, plant spatial networks in drylands organize their linkages fulfilling the structural balance criteria.

**Table 1.**
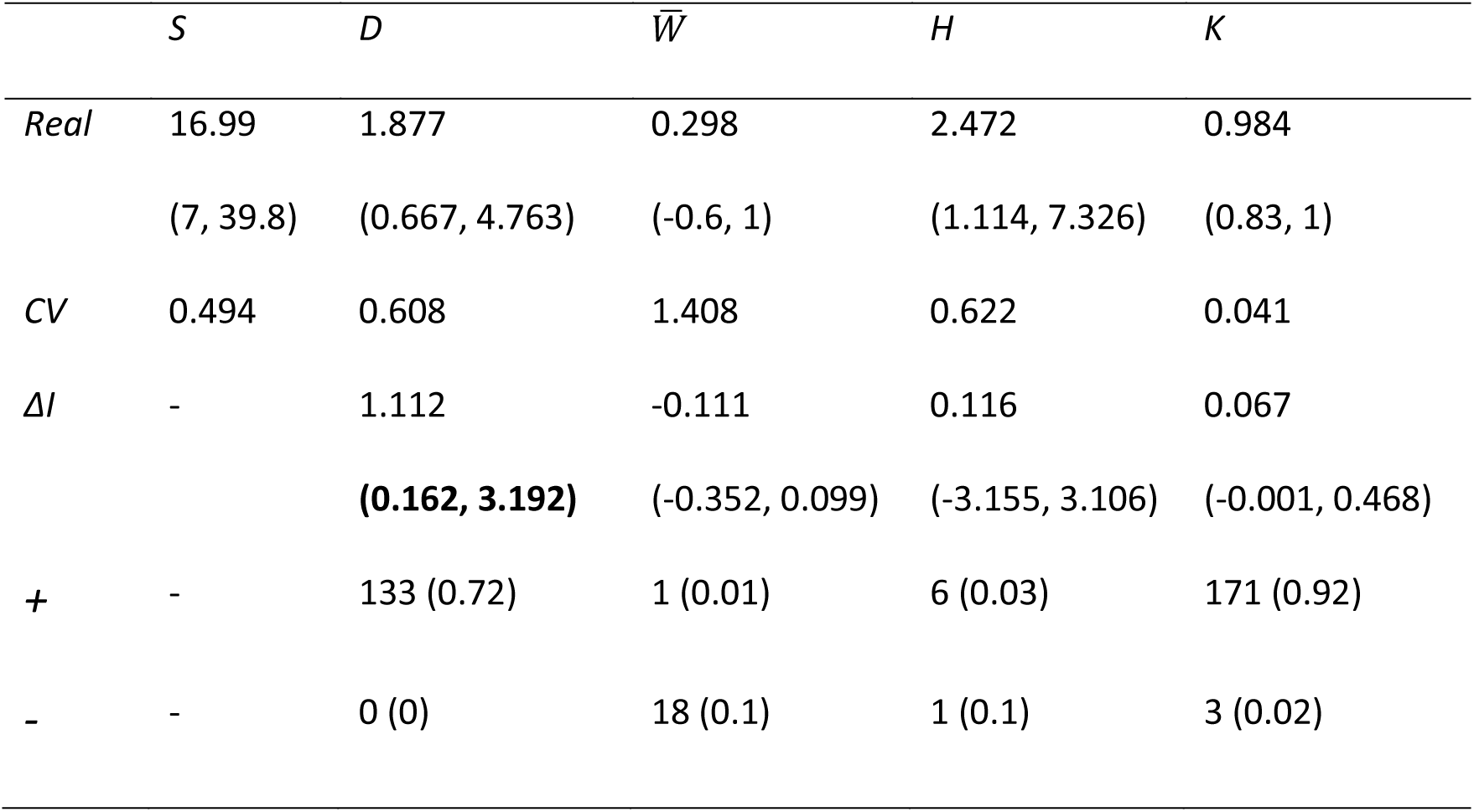
Values of the network indices found in our study sites. *S*, network size; *D*, link density; 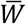, link weight mean; *H*, link weight heterogeneity; *K*, balance. *Real* represents the mean value of the index observed in real networks; *CV* represents the coefficient of variation of *Real*; and *ΔI* represents the mean difference between the index of real networks and the percentile 50 value of their corresponding null model. Values in parentheses represent the 95% confidence interval for the index created using the percentiles 2.5 and 97.5 for study sites; bold values indicate a significant difference between Real and Null values. *+* and − indicate the number of networks which presented significantly higher or lower values for their indices respect the null model (values in parentheses represent the proportion).

| | *S* | *D* | $\bar{W}$ | *H* | *K* |
| --- | --- | --- | --- | --- | --- |
| *Real* | 16.99 | 1.877 | 0.298 | 2.472 | 0.984 |
|  | (7, 39.8) | (0.667, 4.763) | (-0.6, 1) | (1.114, 7.326) | (0.83, 1) |
| *CV* | 0.494 | 0.608 | 1.408 | 0.622 | 0.041 |
| $\Delta I$ | - | 1.112 | -0.111 | 0.116 | 0.067 |
|  |  | **(0.162, 3.192)** | (-0.352, 0.099) | (-3.155, 3.106) | (-0.001, 0.468) |
| $+$ | - | 133 (0.72) | 1 (0.01) | 6 (0.03) | 171 (0.92) |
| $-$ | - | 0 (0) | 18 (0.1) | 1 (0.1) | 3 (0.02) |

**Figure 1.**
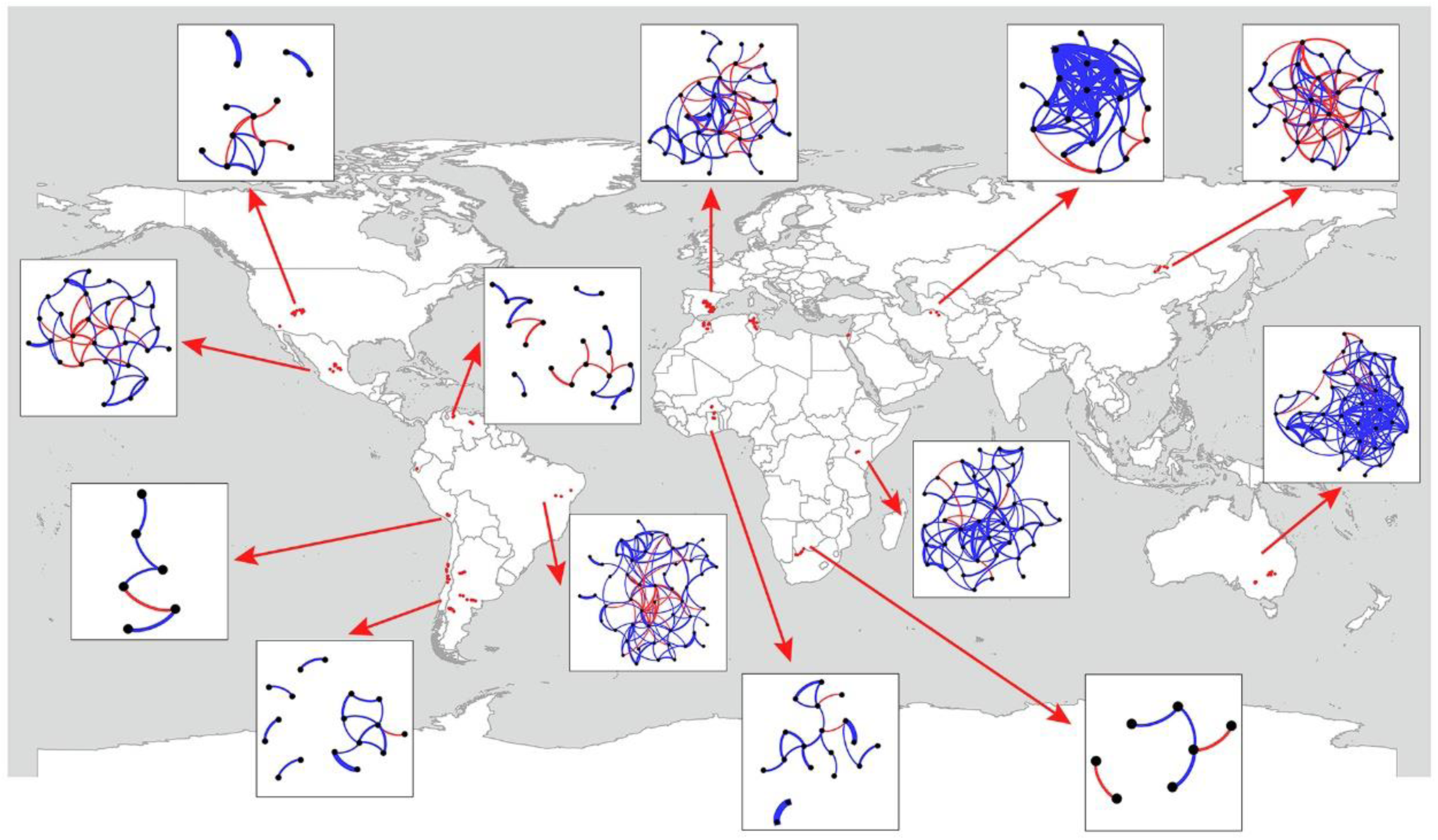
World map showing the locations of all study sites and selected examples of the plant spatial networks found. Blue and red links represent positive and negative interactions, respectively, and link width is proportional to link weight. For simplicity we removed from each network all the species that did not present any link to other species.

The studied plant spatial networks did not organize randomly (Table 1). Specifically, plant communities showed significantly more spatial associations per species (*D*) than expected. Furthermore, 72% of communities presented significantly higher *D* values, and no single community had a lower *D* than expected. These results confirm that plant communities in drylands present a strong spatial structure. Furthermore, 92% of plant communities are closer to the optimal *K* than to the expected value, indicating the prevalence of balanced spatial structures in drylands.

Our structural equation models revealed that the structure of spatial networks significantly affected the richness and evenness of dryland plant communities (Table 2). All network indices had a significant effect on plant community species richness (*SR*), but only *H* and *K* had a significant effect on community evenness (*E*). Furthermore, network variables were the single predictors that had a higher explanatory power on *SR*, while abiotic variables had similar or higher effect than network variables on *E* (Figure 2). Interestingly, the effects of network indices were largely independent from those of other variables, albeit geography and topography had a significant relationship with 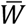 and climate and soil affected *H* (Figures 3 and S2).

**Table 2.** Summary of structural equation models showing the effects of network indices on species richness (*SR*) and evenness (*E*). *D*, link density; 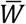, link weight mean; *H*, link weight heterogeneity; *K*, balance; *SR*, community species richness; *E*, community evenness. Bold values indicate a significant direct effect of network index on diversity index. **p < 0.01; ***p < 0.001.

| Network index | Biodiversity index | Path estimate | Standard error | z-value | p-value |
| --- | --- | --- | --- | --- | --- |
| *D* | *SR* | 0.371 | 0.062 | 5.961 | **<0.001\*\*\*** |
|  | *E* | 0.103 | 0.067 | 1.537 | 0.103 |
| $\bar{W}$ | *SR* | 0.227 | 0.07 | 3.253 | **0.001\*\*** |
|  | *E* | 0.014 | 0.069 | 0.207 | 0.836 |
| *H* | *SR* | -0.446 | 0.061 | -7.616 | **<0.001\*\*\*** |
|  | *E* | -0.256 | 0.068 | -3.739 | **<0.001\*\*\*** |
| *K* | *SR* | 0.445 | 0.06 | 7.391 | **<0.001\*\*\*** |
|  | *E* | 0.186 | 0.066 | 2.82 | **0.005\*\*** |

**Figure 2.**
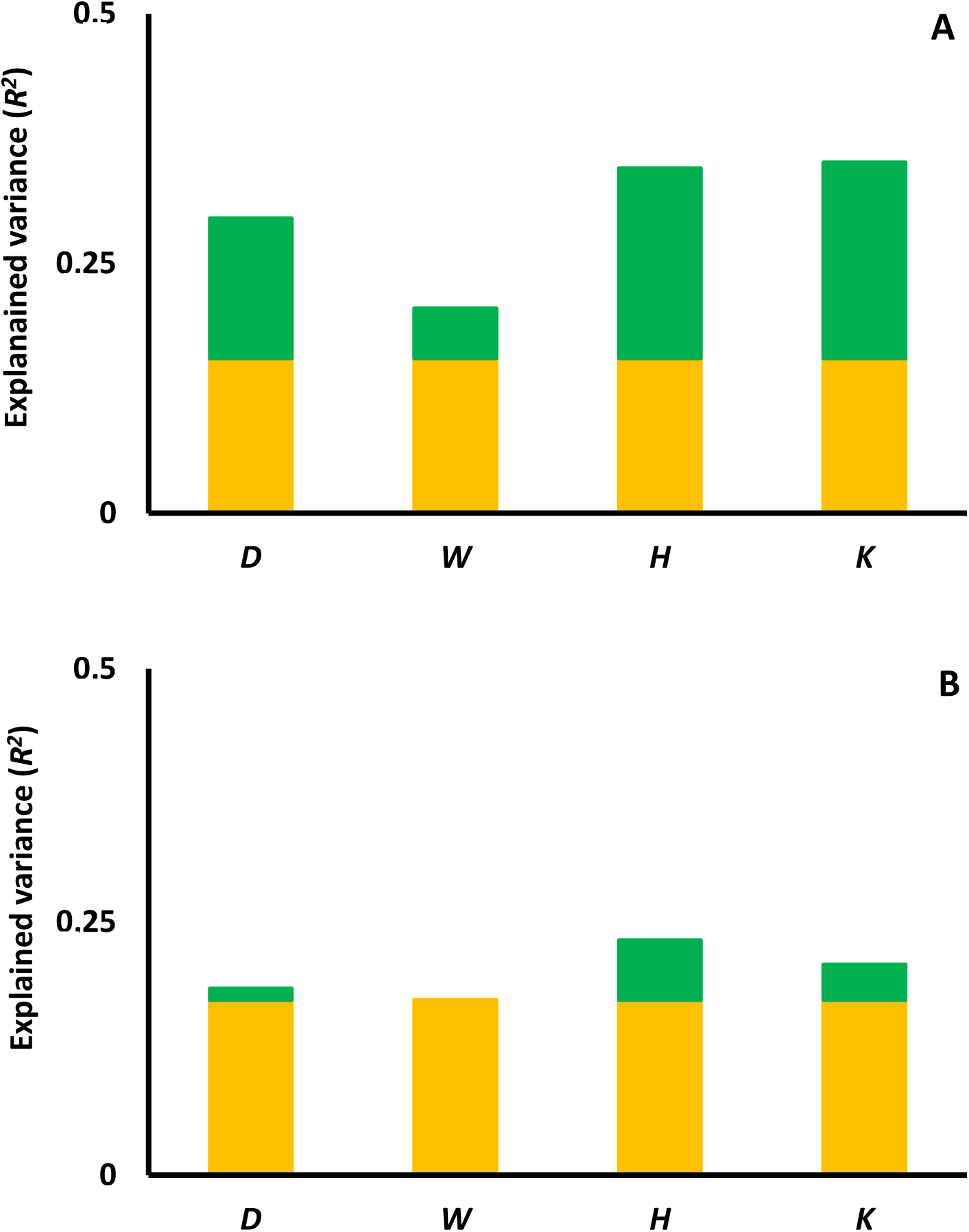
Effects of explanatory variables on species richness (A) and evenness (B). The orange part of the bars represents the explanatory power (*R*^2^) of all abiotic factors together on diversity (both direct and indirect effects); the green part of the bars represents the direct contribution of including each network variable in the structural equation models. *D*, link density; 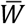, link weight mean; *H*, link weight heterogeneity; *B*, global balance.

**Figure 3.**
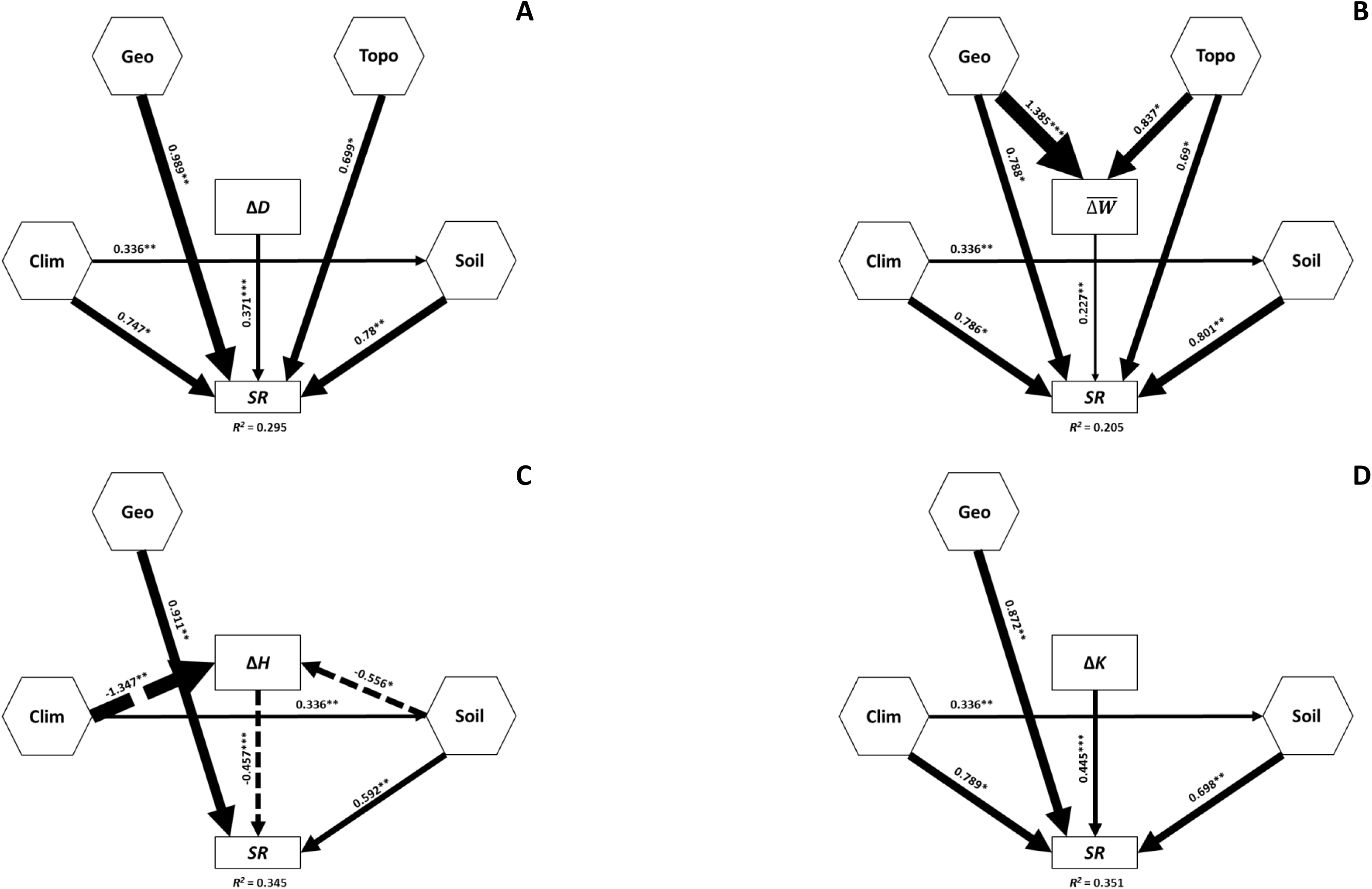
Structural equation models (SEM) describing the effects of abiotic drivers and spatial network indices on plant community diversity. *Geo*, geographical factors; *Topo*, Topographical factors; *Clim*, climatic factors; *Soil*, Soil factors; *SR*, community species richness. Different SEMs represent different network indices: (A) Δ*D*, link density; (B) 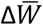, link weight mean; (C) Δ*H*, link weight heterogeneity; and (D) Δ*K*, balance. All network indices are the difference the between real value and the percentile 50 of their respective null model. Numbers adjacent to arrows are indicative of the effect size of the relationship and its significance. Continuous and dashed arrows indicate positive and negative relationships, respectively. *R*^2^ denotes the proportion of variance explained for *SR*. Hexagons are composite variables and squares are observable variables. All models presented a *p*-value > 0.05 for the *χ*^2^. For graphical simplicity, only significant arrows and variables with at least one significant relationship are presented.

## Discussion

Studies at global scales offer unparalleled insights to build generalities in ecology based on the discovery of common patterns and processes operating in a large number of locations and/or ecosystems (Fraser *et al*., 2013). Studies on biotic interactions have often found several network structures that commonly repeat in ecological communities, such as nested and modular patterns (Olesen *et al*., 2007; Thébault & Fontaine, 2010). Some of these structures have been confirmed by global studies conducted on mutualistic systems such as plant-pollinators (Traveset *et al*., 2016), suggesting that biotic interactions at the community level may be structured following general rules. Our results indicate that perennial plant communities present a common network structure, and that network attributes such as high connectivity, link heterogeneity, and balanced structure are positively related to plant diversity in drylands worldwide. Therefore, it is possible that general processes operating in drylands lead to a particular community structure and, as suggested by our results, that this particular structure contributes to increment plant diversity regardless other abiotic factors. Our results constitute, to the best of our knowledge, the first empirical evidence showing the existence of a common network architecture structuring terrestrial plant communities at the global scale, and provide novel evidence of the importance of the network of interactions for the maintenance of biodiversity (Bascompte & Jordano, 2007).

### Plant spatial networks in drylands are highly connected and balanced

Drylands are characterized by particular vegetation patterns, composed by bare soil and vegetation patches (Klausmeier, 1999). Theoretical and empirical results have found that this pattern is the result of hydrological-plant interactions, with bare soil areas acting as ‘sources’ and vegetation patches acting as ‘sinks’ for runoff water after precipitation events (Puigdefabregas *et al*., 1999; Rietkerk *et al*., 2004). Furthermore, empirical and modelling studies have shown a connection between vegetation patchiness and ecosystem processes. For example, Maestre (2006) reported a positive relationship between the spatial pattern of vegetation patches and their water use efficiency, and Berdugo *et al*. (2017) found that this pattern was related to ecosystem multifunctionality (*i.e*. the simultaneous provision of multiple ecosystem functions) in global drylands, being also able to identify the bimodal distribution of multifunctionality observed. However, these studies consider vegetation as a single unity while in general vegetation patches are composed by multiple species that are interacting among them (Tielbörger & Kadmon, 2000), and can respond differently to the same environmental factor (Saiz & Alados, 2011; Pueyo *et al*., 2013).

We found that most plant species presented many spatial associations among them, and that dryland communities could be dominated by spatial aggregation or segregation, as found in many local studies (Fowler, 1986; Soliveres & Maestre, 2014). An interesting pattern observed is that, regardless of the dominant spatial pattern found at each site, vegetation patches in drylands seem to organize following the structural balance criteria worldwide. Thus, within a given plant community, plant species that aggregate within the same patches do not appear in patches formed by other species (Saiz *et al*., 2016). In drylands, species responsible of patch formation facilitate the establishment of seedlings under their canopies, but when seedlings become adults, these interaction may change to competition (Tielbörger & Kadmon, 2000). Particularly, this change is more common between phylogenetically related species which share niche requirements, resulting in communities where plant species tend to interact negatively with close relative species and positive with a subset of the distant relatives (Verdú *et al*., 2010). In our case, as we only consider mature plant communities, we could expect that the different blocks observed within our networks represent the different types of vegetation patches present in the community (Saiz *et al*., 2014), and that the balanced organization is the result of niche processes among species (*e.g*. promotion of competition between closely related species and facilitation between distant ones, Verdú & Valiente-Banuet, 2011). Furthermore, although not specifically tested in ecology so far, studies on signed networks suggest that the balance criterion would promote the resilience of the network (*e.g*. reducing the disturbances within the network, Cartwright & Harary, 1956; or increasing the adaptability of the system, Ilany *et al*., 2013). Hence, our results suggest that the balanced spatial structures observed could increase the resilience of dryland plant communities worldwide.

Substantial research efforts have been developed to understand the links between network structure and community robustness. For example, the nested structure of mutualistic networks makes the community robust to the random extinction of species (Memmott *et al*., 2004), while modularity is linked to higher resilience in trophic networks (Thébault & Fontaine, 2010). A recent study by (Gao *et al*., 2016) posits that both network resilience against the random extinction of nodes and changes in the number and strength of links increase with higher link density and more heterogeneous link strength distributions. The plant spatial association networks studied here present one of these conditions, high link density, while we did not found a significant difference for link strength heterogeneity between real networks and the null model. This could be explained by the high dominance of links with low weight, which is the typical case for ecological networks and it has been suggested to play an essential role maintaining the persistence in food webs (McCann *et al*., 1998). Thus, our results suggest that plant spatial association networks present a structure which enhances community resilience against perturbations affecting both species and interactions among them.

### The structure of plant spatial networks promotes species diversity

We found a significant effect of network indices on plant species richness and evenness in drylands worldwide. All indices linked to network resilience, such as *D* and *H*, had positive effects on the diversity metrics studied (higher link density and link weight heterogeneity values where linked to a higher plant diversity), and the positive effect of *K* on diversity supports the idea that balanced spatial structures increase the resilience of plant communities. On the other hand, and contrary to our expectations, we did not find any effect of the importance of positive interactions on plant diversity. A possible explanation is that the importance of positive and negative interactions alone (*sensu* Brooker *et al*., 2005) does not suffice to explain the coexistence of diverse species in a community, and thus is imperative to explore also the structure of these interactions (Soliveres *et al*., 2015b). For example, it has been proposed that coexistence of more species can be enhanced when competitive interactions are not hierarchical, but are intransitive (Wootton, 2001), something that has been found in a previous analyses of our global database (Soliveres *et al*., 2015a). Therefore, our results encourage the use of approaches as networks in plant communities, which not only account for the importance of biotic interactions but also for their structure in the community, and are able to consider simultaneously different types of interactions as facilitation and competition.

Additionally, we found that the effects of network structure on the diversity of plant communities are largely independent from those of abiotic factors. Previous studies conducted in arid environments have suggested that nested network structures of facilitative interactions help to preserve the diversity of plant communities (Verdú *et al*., 2008). Furthermore, a positive relationship between the spatial organization of vegetation patches and plant species richness has also been found (with more patchy communities associated to higher number of species; Maestre, 2006; Pueyo *et al*., 2013). Our results represent a step forward, as the network approach used considers both facilitation and competition, and show that the structure of both facilitative and competitive interactions have a direct effect in plant community diversity. On the other hand, recent studies have found that attributes such as plant species richness are positively related to multifunctionality in drylands (Maestre *et al*., 2012b), which is also affected by other community attributes such as species composition and spatial pattern (Maestre *et al*., 2012a; Berdugo *et al*., 2017). Therefore, it is very likely that the structure of plant interaction networks is not only responsible for the resilience of the community against the extinction of species and interactions, but also plays a key role in the maintenance of ecosystem functions against future environmental changes.

### Concluding remarks

The analysis of the plant spatial association networks revealed new insights on the structure of plant communities in drylands worldwide. These communities showed common patterns but, in contrast to previous studies focused on local communities and positive interactions (Verdú *et al*., 2008), we found that these patterns apply worldwide to plant communities including both positive and negative interactions. The structure of the networks studied showed a high density of connections between species and followed a balanced criterion, properties that are related with the resilience of the communities against disturbances (Gao *et al*., 2016). Furthermore, networks with dense and heterogeneous connections and balanced structures presented higher plant diversity, which supported the idea of these network structures promoting the coexistence of larger number of species. Finally, the independence of these properties from abiotic factors and from the dominance of positive or negative links revealed the need to take into account not only the importance of biotic interactions but also their structure when studying drivers of dryland vegetation assembly. Although challenging, global field studies provide the framework to find common patterns in natural communities and to advance in our understanding of ecosystem processes. Our results highlight the importance of system level approaches to explain the diversity of plant species, a major driver of ecosystem functioning, and to identify the structure of communities that is likely to provide the highest resilience against disturbances, in drylands worldwide.

## Acknowledgements

We thank all the members of the EPES-BIOCOM network for the collection of field data and all the members of the Maestre lab for their help with data organization and management, and for their comments and suggestions on early stages of the manuscript. This work was funded by the European Research Council under the European Community’s Seventh Framework Programme (FP7/2007-2013)/ERC Grant agreement 242658 (BIOCOM). FTM and HS are supported by the European Research Council (ERC Grant agreement 647038 [BIODESERT]); and JGG acknowledges financial support from MINECO (through projects FIS2015-71582-C2 and FIS2014-55867-P) and from the Departamento de Industria e Innovación del Gobierno de Aragón y Fondo Social Europeo (FENOL group E-19).

## Supplementary material

S1. Structural equation model used to evaluate the effect of network indices on plant community diversity and explanation of the variables.

S2. Structural equation models (SEM) describing the effects of abiotic drivers and spatial network indices on plant community evenness.

## Data availability statements

All the materials, raw data, and protocols used in the article are available upon request and without any restriction, and will be published in an online repository (figshare) upon the acceptance of the article (currently available as a private link https://figshare.com/s/6ae331fa92446433d8a4).

## Biosketch

Hugo Saiz is a community ecologists specially interested in disentangling the mechanisms that structure plant communities and allow the coexistence and persistence of high diverse ecosystems. Fernando Maestre is a drylands ecologist and his research uses an integrative approach combining a wide variety of tools, biotic communities and scales to understand the functioning of semiarid ecosystems. Jesús Gómez-Gardeñes and Juan Pablo Borda are physicists working on the dynamic of complex systems including both theoretical approaches and its application to real systems ranging from car displacements to ecological systems.

